# Multiplatform Integrative Analyses of Immunosuppressive Signatures in Cortisol-secreting Adrenocortical Carcinoma

**DOI:** 10.1101/2021.01.14.426712

**Authors:** Jordan J. Baechle, David N. Hanna, Sekhar R. Konjeti, Jeffrey C. Rathmell, W. Kimryn Rathmell, Naira Baregamian

## Abstract

Adrenocortical carcinoma (ACC) is a rare but highly aggressive malignancy and nearly half of ACC tumors have been shown to overproduce and secrete adrenal steroids. Excess cortisol secretion, in particular, has been associated with poor prognosis among ACC patients. Furthermore, recent immunotherapy clinical trials demonstrated significant immunoresistance among cortisol-secreting ACC (CS-ACC) patients when compared to their non-Cortisol-secreting (nonCS-ACC) counterparts. The immunosuppressive role of excess glucocorticoid therapies and secretion is well established, however, the impact of the cortisol hypersecretion on ACC tumor microenvironment (TME), immune expression profiles, and immune cell responses remain largely undefined. In this study, we characterized the TME of ACC patients and compared the immunogenomic profiles of nonCS-ACC and CS-ACC tumors to assess the impact of differentially expressed genes (DEGs) related to immune processes on patient prognosis. Comprehensive multiplatform immunogenomic computational analyses of ACC tumors deciphered an immunosuppressive expression profile with a direct impact on patient survival. We identified several primary immunogenomic prognostic indicators and potential targets within the tumor immune landscape of CS-ACC that define a distinct TME and provide additional insight into the understanding of potential contributory mechanisms underlying failure of initial immunotherapeutic trials and poor prognosis of patients with CS-ACC.

## Introduction

Adrenocortical carcinoma (ACC) is among the rarest and most aggressive cancers. Although the current prognostication of patients with ACC primarily hinges on the presence or absence of metastases and tumor resectability, over a third of patients present with an advanced, esectable disease (1-6). Patients with the fully resectable (R_0_) disease have a reported 5-year survival rate of approximately 50%, whereas patients with the unresectable disease have a 5-year survival rate near 0% and a median survival of fewer than 12 months (4,7-8). Aside from the advent of mitotane therapy in the treatment of ACC in 1959, there has been little improvement in overall mortality over the past several decades (9-10). Despite limited therapeutic options for patients with unresectable ACC, several immunotherapies are currently under evaluation (11-12).

Nearly half of patients presenting with ACC have been shown to exhibit steroid hormone hypersecretion with excess cortisol secretion being the most predominant hormone, and often considered an indicator of poor prognosis (13-14). Glucocorticoids, including cortisol, are small lipid hormones produced by the adrenal glands that exert their effects through glucocorticoid receptors modulating tumor expression to perform a variety of functions, including arresting immune cell growth and maturation, inhibiting activation signaling, and inducing apoptosis in lymphocytes (15-16). Glucocorticoids have proven so effective in this role that they are the cornerstone of treatment for many hypersensitivity immune reactions and autoimmune diseases (17-19). The immunosuppressive effects of excess glucocorticoid therapy and hypersecretion, however, have also been shown to hinder the immune system’s capacity to ward off infections and malignancy and have been associated with a variety of other effects, including muscle wasting, osteoporosis, and metabolic derangements (20-22). The most recent immunotherapy clinical trials revealed a significant pattern of immune resistance among cortisol-secreting ACC (CS-ACC) tumors, with higher rates of immunotherapeutic failure among CS-ACC patients compared to the patients with nonCS-ACC (23-26).

In this study, we define the suppression of immune response signatures within the tumor-infiltrating immune cell (TIIC) landscape of CS-ACC using comprehensive multiplatform computational immunogenomic analytic approaches. Furthermore, we explore the clinical relevance of these immunogenomic findings to further decipher the immune resistant behavior of CS-ACC tumors for future targeted immunotherapeutic approaches. We performed a comparative analysis of the TME immunogenomic landscapes and TIIC profiles of CS- and nonCS-ACC tumors and analyzed the prognostic value of differentially expressed genes and infiltrating immune cell estimations as well as intercorrelated relationships. Multiplatform analyses identified primary immunogenomic prognostic indicators and therapeutic targets that may advance the overall understanding of the role of TIICs in ACC biology and the influence of cortisol secretion in ACC TME.

## Results

We identified 79 individuals with ACC tumors and 33 (42%) had CS-ACC and 46 (58%) were nonCS-ACC tumors (Table 1). The groups were similar in age at diagnosis (p=0.553), race (p=0.187), tumor stage T (p=0.581), nodal status N (p=0.131), metastasis M (p=0.769) and clinical stage (p=0.234). The CS-ACC group was significantly female-predominant compared to the nonCS-ACC group (81.8 vs. 46.3%, p=0.004). CS- and nonCS-ACC tumors demonstrated similar fractions of genome alteration (p=0.599), mutation count (p=0.139), mitotic rate (p=0.509), tumor necrosis (p=0.498), Weiss Score (p=0.848) (22), and rates of vascular invasion (p=0.401). CS-ACC tumors demonstrated significantly elevated mitotic count compared to nonCS-ACC tumors (9.0 vs. 5.0, p=0.030). Both groups reported similar resection margins (p=0.616) and underwent similar rates of unspecified neoadjuvant (p=0.458) and adjuvant (p=0.125) therapy, as well as adjuvant radiation therapy (p=0.401). CS-ACC patients were significantly more likely to receive adjuvant mitotane therapy (78.8 vs. 53.8%, p=0.037) and experienced higher rates of ACC recurrence (67.7 vs 28.6%, p=0.003). Demographic, clinical, and pathologic features of the study cohort by cortisol secretion are further summarized in Table 1.

**Table 1.** Patient Demographic, Tumor Pathology, and Treatment Parameters of Adrenocortical Carcinoma (ACC)*.

|  |  | NonCS-ACC<br>N=39 | CS-ACC<br>N=33 | p-value |
| --- | --- | --- | --- | --- |
| **Age (years) -Median [IQR]** |  | 50.0 [36.0;57.0] | 47.0 [34.0;61.0] | 0.553 |
| **Sex** | Female | 18 (46.2%) | 27 (81.8%) | **0.004** |
|  | Male | 21 (53.8%) | 6 (18.2%) |  |
| **Race** | White | 31 (79.5%) | 30 (90.9%) | 0.187 |
|  | Asian | 1 (2.56%) | 0 (0.00%) |  |
|  | Black | 0 (0.00%) | 1 (3.03%) |  |
|  | Unknown | 7 (17.9%) | 2 (6.06%) |  |
| **Clinical Stage** | Stage I | 5 (12.8%) | 3 (9.09%) | 0.234 |
|  | Stage II | 21 (53.8%) | 12 (36.4%) |  |
|  | Stage III | 5 (12.8%) | 10 (30.3%) |  |
|  | Stage IV | 7 (17.9%) | 8 (24.2%) |  |
|  | Unknown | 1 (2.56%) | 0 (0.00%) |  |
| **T Stage** | T1 | 5 (12.8%) | 3 (9.09%) | 0.581 |
|  | T2 | 22 (56.4%) | 15 (45.5%) |  |
|  | T3 | 3 (7.69%) | 5 (15.2%) |  |
|  | T4 | 8 (20.5%) | 10 (30.3%) |  |
|  | Unknown | 1 (2.56%) | 0 (0.00%) |  |
| **N Stage** | N0 | 36 (92.3%) | 27 (81.8%) | 0.131 |
|  | N1 | 2 (5.13%) | 6 (18.2%) |  |
|  | Unknown | 1 (2.56%) | 0 (0.00%) |  |
| **M Stage** | M0 | 31 (79.5%) | 25 (75.8%) | 0.769 |
|  | M1 | 7 (17.9%) | 8 (24.2%) |  |
|  | Unknown | 1 (2.56%) | 0 (0.00%) |  |
| **ACC Type** | Common Type | 35 (89.7%) | 33 (100%) | 0.245 |
|  | Oncocytic Type | 3 (7.69%) | 0 (0.00%) |  |
|  | Myxoid Type | 1 (2.56%) | 0 (0.00%) |  |
| **Excess Hormone Secretion** | Cortisol | 0 (0.00%) | 16 (48.5%) | <0.001 |
|  | Androgen + Cortisol | 0 (0.00%) | 16 (48.5%) |  |
|  | Androgen | 8 (20.5%) | 0 (0.00%) |  |
|  | Estrogen | 2 (5.13%) | 0 (0.00%) |  |
|  | Mineralocorticoids | 3 (7.69%) | 0 (0.00%) |  |
|  | Mineralocorticoid + Cortisol | 0 (0.00%) | 1 (3.03%) |  |
|  | None | 26 (66.7%) | 0 (0.00%) |  |
| **Laterality** | Left | 22 (56.4%) | 18 (54.5%) | 1.00 |
|  | Right | 17 (43.6%) | 15 (45.5%) |  |
| **Fraction of Genome Altered -Median [IQR]** |  | 0.65 [0.43;0.89] | 0.48 [0.32;0.91] | 0.599 |
| **Mutation Count -Median [IQR]** |  | 82.5 [69.0;106] | 94.5 [78.8;118] | 0.139 |
| **Mitoses Count -Median [IQR]** |  | 5.00 [3.00;11.5] | 9.00 [5.75;36.0] | **0.030** |
| **Mitotic Rate** | < 5/50 HPF | 18 (46.2%) | 11 (33.3%) | 0.509 |
|  | > 5/50 HPF | 17 (43.6%) | 19 (57.6%) |  |
|  | Unknown | 4 (10.3%) | 3 (9.09%) |  |
| **Tumor Necrosis** | Absent | 11 (28.2%) | 6 (18.2%) | 0.498 |
|  | Present | 27 (69.2%) | 25 (75.8%) |  |
| **Weiss Score -Median [IQR]** |  | 5.00 [3.50;7.00] | 6.00 [4.00;8.00] | 0.848 |
| **Venous Invasion** | Absent | 23 (59.0%) | 14 (42.4%) | 0.401 |
|  | Present | 13 (33.3%) | 16 (48.5%) |  |
|  | Unknown | 3 (7.69%) | 3 (9.09%) |  |
| **Neoadjuvant Therapy** | No | 39 (100%) | 32 (97.0%) | 0.458 |
|  | Yes | 0 (0.00%) | 1 (3.03%) |  |
| **Resection Margin** | R0 | 28 (71.8%) | 21 (63.6%) | 0.616 |
|  | R1 | 3 (7.69%) | 3 (9.09%) |  |
|  | R2 | 4 (10.3%) | 5 (15.2%) |  |
|  | RX | 2 (5.13%) | 4 (12.1%) |  |
|  | Unknown | 2 (5.13%) | 0 (0.00%) |  |
| **Adjuvant Therapy** | No | 14 (35.9%) | 8 (24.2%) | 0.125 |
|  | Yes | 22 (56.4%) | 25 (75.8%) |  |
|  | Unknown | 3 (7.69%) | 0 (0.00%) |  |
| **Adjuvant Mitotane Therapy** | No | 15 (38.5%) | 7 (21.2%) | **0.037** |
|  | Yes | 21 (53.8%) | 26 (78.8%) |  |
|  | Unknown | 3 (7.69%) | 0 (0.00%) |  |
| **Adjuvant Radiation Therapy** | No | 30 (76.9%) | 23 (69.7%) | 0.401 |
|  | Yes | 6 (15.4%) | 9 (27.3%) |  |
|  | Unknown | 3 (7.69%) | 1 (3.03%) |  |
| New Neoplasm<br>Following Initial<br>Treatment | No | 25 (71.4%) | 10 (32.3%) | 0.003 |
|  | Yes | 10 (28.6%) | 21 (67.7%) |  |

**\*Abbreviations:** CS-ACC = cortisol-secreting adrenocortical carcinoma; nonCS-ACC = non-cortisol-secreting adrenocortical carcinoma; IQR = interquartile range; Age = age at diagnosis; Unknown = unreported data; HPF = high-power fields; RX = indeterminate margins

Univariate Cox regression and Kaplan-Meier analysis examined the impact of excess cortisol secretion on overall survival (OS) and disease-free survival (DFS). Cortisol secretion was significantly associated with shortened OS (hazard ratio [HR] 2.28; 95% confidence interval [CI] 1.08 – 4.83, p=0.031) and DFS (HR 2.42; 95% CI 1.26 – 4.68, p<0.01). The 5-year OS was 68% for nonCS-ACCs and 50.1% for CS-ACCs (p=0.026) (Figure 1A) and the 5-year DFS was 61.2% and 29.2% for nonCS- and CS-ACC tumors (p=0.006), respectively (Figure 1B). The poor prognosis related to CS-ACC despite similar patient demographics, tumor pathology, and treatment protocols commonly associated with survival (patient age, cancer stage, Weiss Score, neoadjuvant/adjuvant therapy) is suggestive of a possible direct impact of cortisol secretion on ACC biology or immune opposition underlying patient OS and DFS.

**Figure 1.**
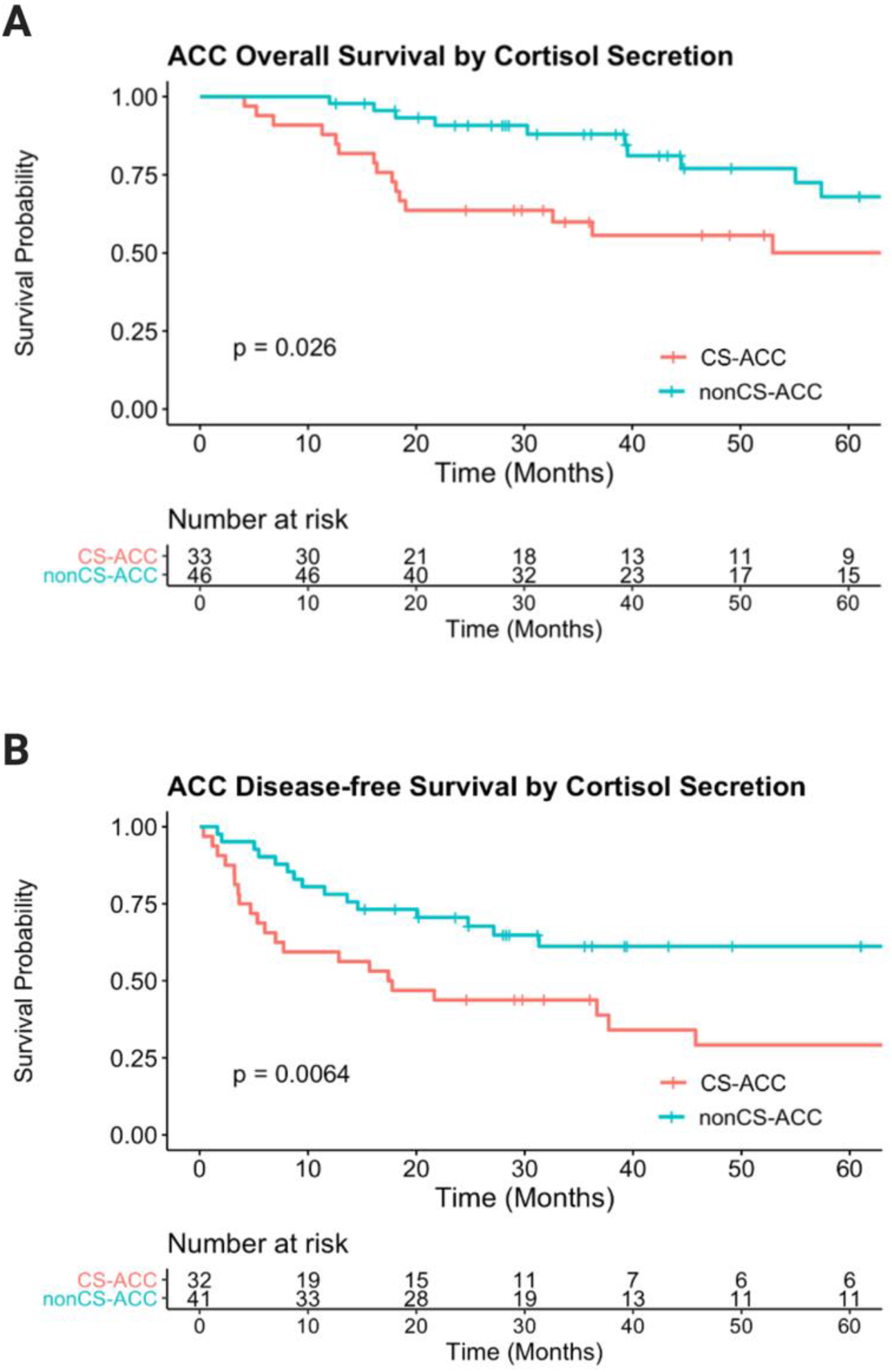
Patient Overall Survival and Disease-Free Survival Outcomes by Cortisol Secretion in Adrenocortical Carcinoma (ACC). (**A)**. Overall survival extrapolated from The Cancer Genome Atlas (TCGA) cohort of patients with adrenocortical carcinoma (N=79) is stratified by cortisol secretion. ACC, adrenocortical carcinoma; CS-ACC, cortisol-secreting ACC; nonCS-ACC, non-cortisol-secreting ACC. (**B)**. Disease-free survival is stratified by cortisol secretion in patients with ACC.

Analysis of all genes (n>19,000) with quantified mRNA expression in The Cancer Genome Atlas (TCGA) database demonstrated 1,612 differentially expressed genes (DEGs) between CS- and nonCS-ACC tumors (p<0.05 and q<0.05) (14). Of these DEGs, 1,021 were classifiable using Panther Genomic Classification. Forty-five (4%) genes were identified to be directly related to immunological processes (Figure 2A). On subcategorization, the identified immune genes were primarily involved in immune response and leukocyte activation and maturation. The observed differences in the distribution of immunological processes associated with each gene expression and unique mRNA expression profiles of these 45 immune DEGs further defined distinct profile differences between CS- and nonCS-ACC tumors (Figure 2, B and C). Forty-four (98%) of the immune DEGs identified had shown decreased expression levels in CS-ACC compared to nonCS-ACC tumors. Notably, *CCRL2* uniquely showed elevated mRNA expression levels within CS-ACC TME compared to nonCS-ACC. All immune DEGs showed positive mRNA expression intercorrelation with one another (r^2^≥0.00) aside from *CCRL2. CCRL2* mRNA expression was negatively associated with that of several genes, including *CCR6* (r^2^=-0.37), JAK3 (r^2^=-0.30), NKAP (r^2^=-0.26), SIRPA (r^2^=-0.31), and TLR5 (r^2^=-0.27) (Figure 2D).

**Figure 2.**
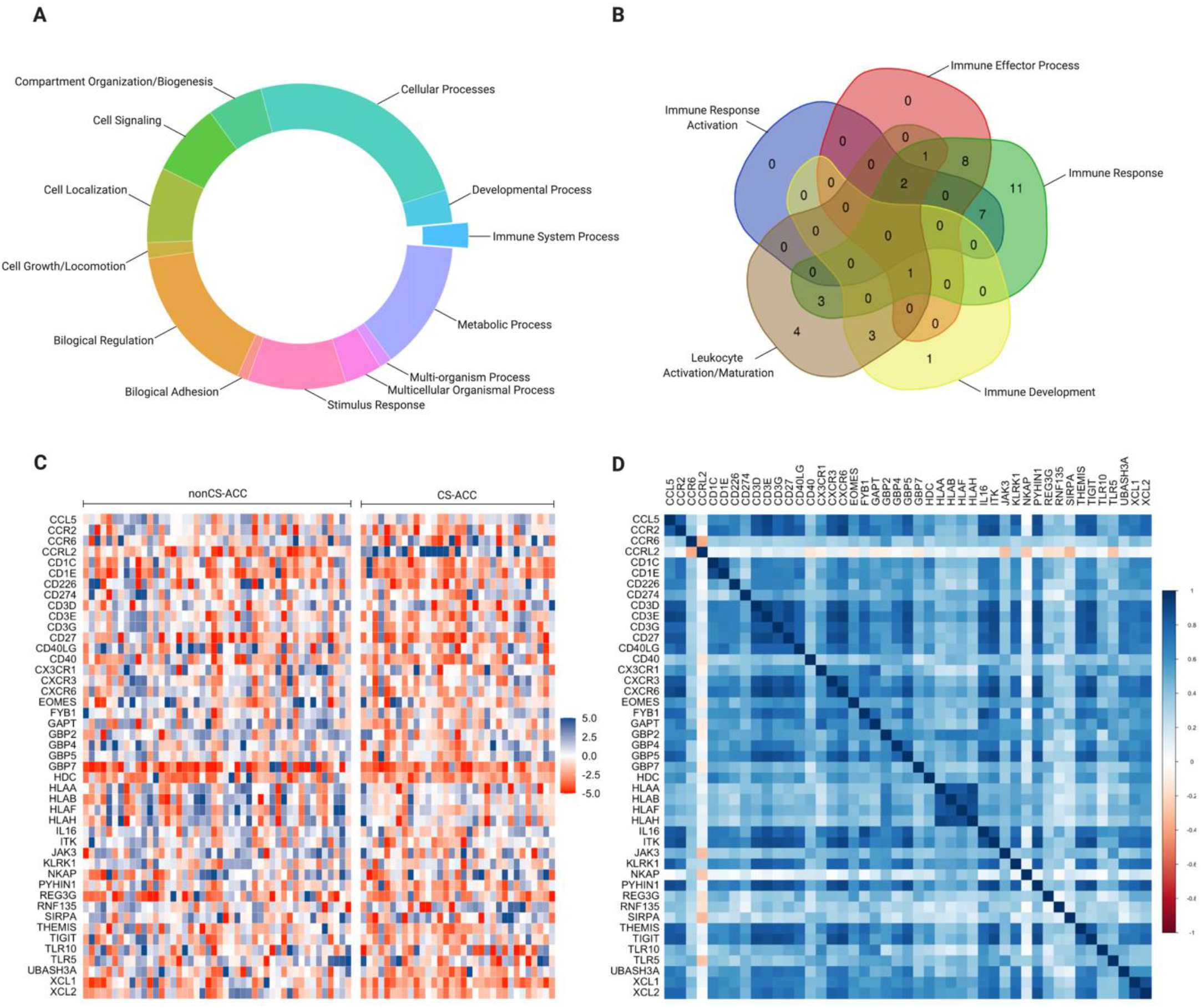
Differential Gene mRNA Expression (DEGs) of Adrenocortical Carcinoma Stratified by Cortisol Secretion. (**A)**. Categorization of all differentially expressed genes (DEGs). (**B)**. Subcategorization of DEGs directly involved in immunological processes. (**C)**. Heatmap of differential gene expression between CS-ACC and nonCS-ACC. (**D)**. Heatmap of relationships between deferentially expressed immune genes.

The immunogenomic TME deconvolution using a computational approach of *CIBERSORTx* analytical platform elucidated a distinct TIIC landscape among CS-ACC compared to nonCS-ACC based on both the proportion and the absolute TIIC estimations (Figure 3, A, B, and C). The average proportional TIIC distribution of CS- and nonCS-ACC (Figure 3A) and the proportional and absolute TIIC estimations were stratified by cortisol secretion to reveal a differential pattern of expression (Figure 3, B and C). The schematic diagram (Figure 4) comprehensively summarizes our global analysis and tumor-infiltrating immune cell profiles of CS-ACC.

**Figure 3.**
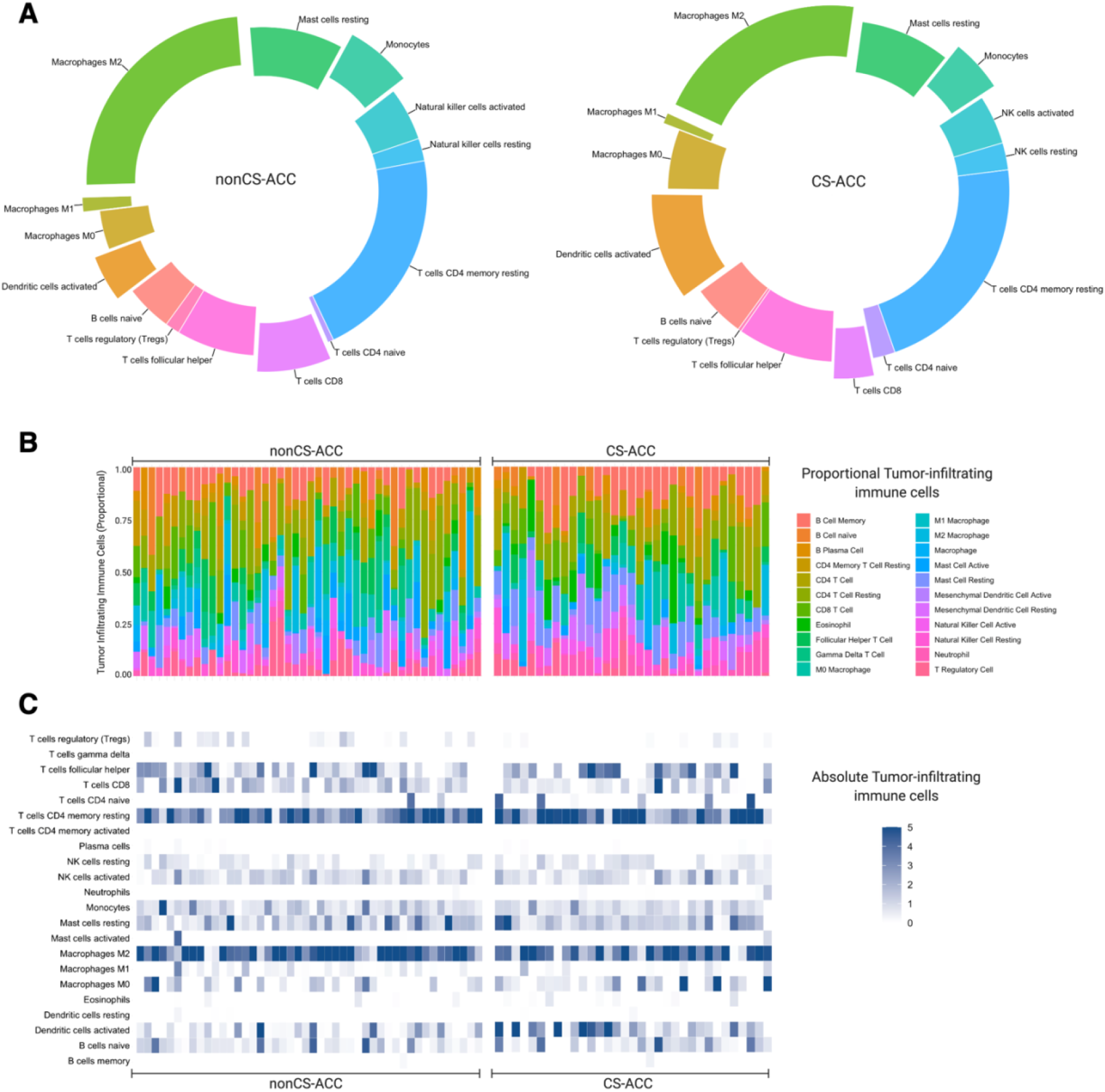
Tumor-Infiltrating Immune Cell (TIIC) Profiles of Adrenocortical Carcinoma (ACC). **(A)**. An average proportional immune cell infiltration stratified by cortisol secretion in ACC tumors. (**B)**. Individual patient proportional relative values of immune cell infiltration stratified by cortisol secretion of ACC tumors. (**C)**. Normalized absolute immune cell infiltration estimates stratified by cortisol secretion of ACC. *CIBERSORTx* computational analytical platform was used to express the estimated tumor-infiltrating immune subsets.

**Figure 4.**
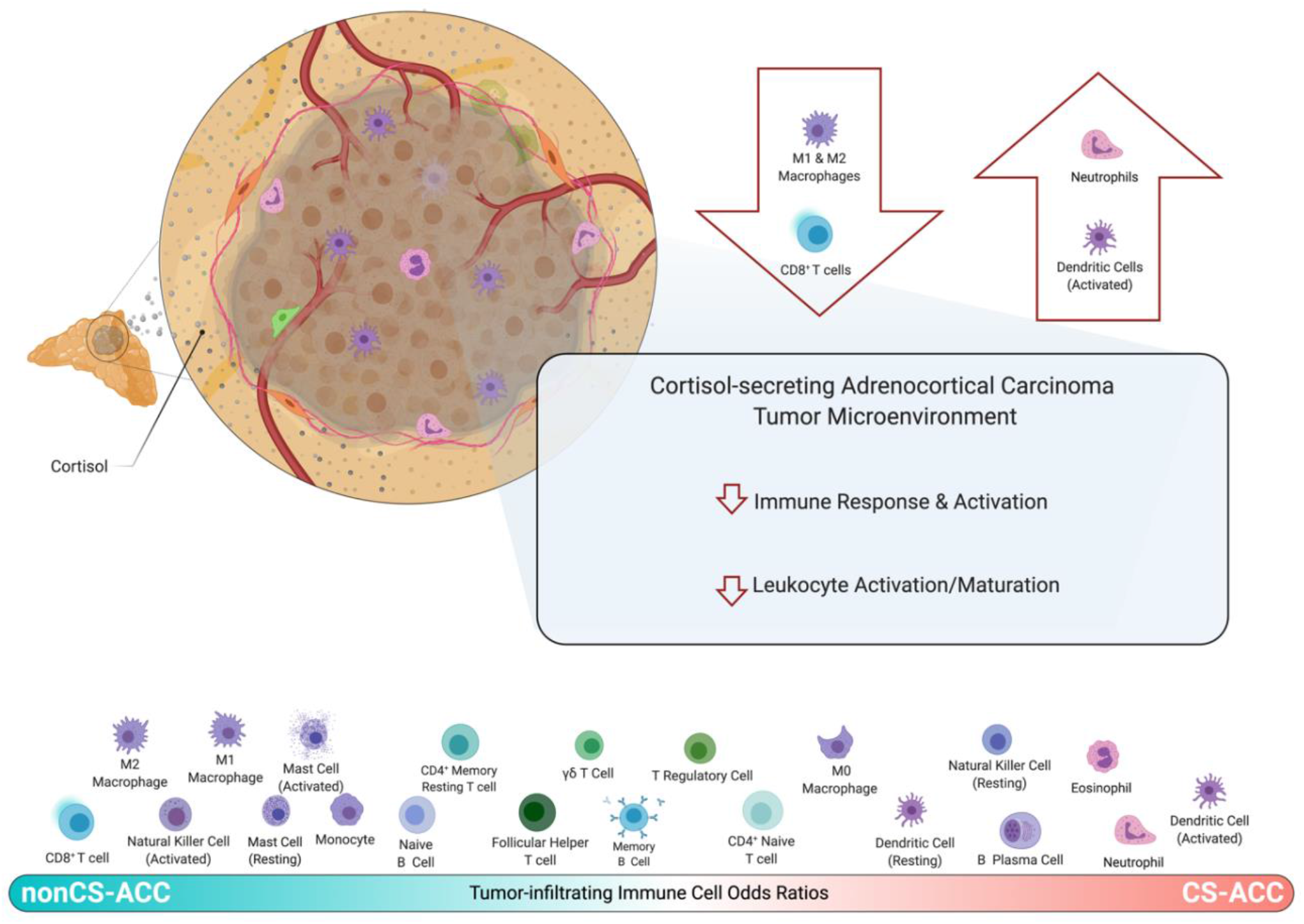
Immune Landscape of Cortisol-Secreting Adrenocortical Carcinoma (CS-ACC) Tumor Microenvironment (TME). Schematic comparison of estimated tumor immune cell type infiltration odds ratios of CS-ACC and nonCS-ACC TME.

A comparison of absolute TIIC populations according to cortisol secretion in the CS-ACC tumors demonstrated significant depletion of CD8^+^ T cells (p=0.017), monocytes (p=0.040), as well as M1 (p=0.048) and M2 macrophages (p=0.030), and increased infiltration of activated dendritic cells (DC_a_) (p=0.016) (Figure 5A). Of these five immune cell types found to be differentially infiltrated in CS- and nonCS-ACC TMEs, four were found to be significant prognostic indicators of OS and/or DFS (p<0.05). CD8^+^ T cell infiltration was associated with improved DFS (HR 0.01, 95% CI 0.00 – 0.53, p=0.023). The presence of monocytes in the TME was shown to be associated with improved OS (HR 0.01, 95% CI 0.00 – 0.53, p=0.008). Tumor infiltration of M2 macrophage was associated with improved OS (HR 0.04, 95% CI 0.01 – 0.37, p=0.004) and DFS (HR 0.03, 95% CI 0.00 – 0.19, p<0.001). Increased immunosuppressive DC_a_ infiltration was strongly associated with poor OS (HR 27.9, 95% CI 2.01 – 88.8, p=0.013) and DFS (HR 78.9, 95% CI 7.51 – 829, p<0.001) The prognostic value of each differentially infiltrated tumor immune cell type is depicted in Figure 5B.

**Figure 5.**
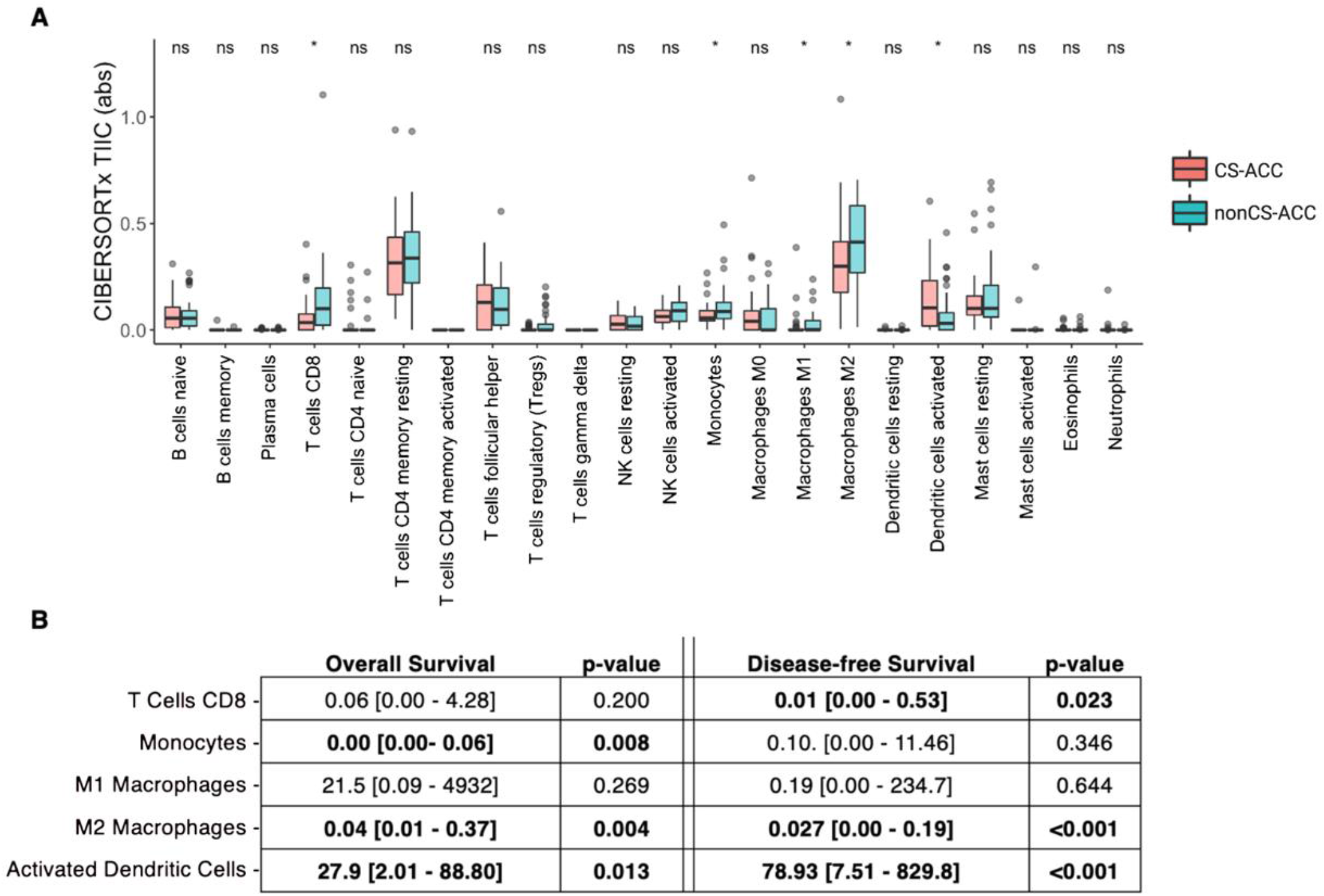
Differential Tumor Immune Cell Infiltration by Cortisol Secretion and Associated Patient Outcomes in Adrenocortical Carcinoma (ACC). **(A)**. Normalized absolute value of tumor infiltration by immune cell types estimated by *CIBERSORTx* in ACC tumors and stratified into subgroups by cortisol secretion, abs = absolute arbitrary units; ns = p-value ≥0.05; * = p-value <0.05. (**B)**. Impact of differentially expressed TIIC subtypes (CD8, Monocytes, M1 and M2 macrophages, activated Dendritic Cells) on Overall (OS) and Disease-Free Survival (DFS). Regression analysis, expressed as univariate Cox regression hazard ratio (HR) and 95% Confidence interval (95% CI): HR [lower 95% CI – higher 95% CI], **bold** = p-value <0.05.

Of the identified 1,612 differentially expressed genes (DEGs) between CS- and nonCS-ACC tumors, 45 were found to be related to immunological processes. Of the 45 immune DEG hits, the mRNA expression of 19 (42%) genes emerged as significant prognostic indicators with positive associations for both OS and DFS in CS-ACC (HR >1.00, OS and DFS, p<0.05). These genes were *CCR2, CCL6, CD1C, CD1E, CD40LG, CD40, CXCR6, EOMES, GAPT, GBP2, HLAA, HLAB, HLAF, HLAH, IL16, JAK3, NKAP, REG3G, SIRPA*. These genes with mRNA expression significant for OS and DFS prognostication can be grouped into several subcategories according to immune function, including chemokine and cytokine signaling (*CCR2, CCL6, CXCR6*), macrophage signaling (*IL16, GBP2, REG3G*), leukocyte antigen proteins, (*HLA-A, B, F, H*), T cell signaling (*CDE1, CD1C, EOMES, NKAP*), B cell signaling (*CD40LG, CD40, GAPT*), dendritic cell signaling (SIRPA), and global immune development and response (*JAK3*) (Figure 6, A, B and C). The comparative distribution of DEGs by cortisol secretion and the impact of DEGs on OS and DFS are summarized in Supplemental Table 1. Seventeen genes were positively associated with DFS and included *CCL5, CD3E, CD27, CXCR3, FYB1, GBP5, GBP7, HDC, IKT, KLRK1, PYHIN1, RNF135, THEMIS, TIGIT, TLR10, XCL1*, and *XCL2* (Figure 6C). Functionally, these genes can be grouped into chemotactic signaling (*CCL5, CXCR3, XCL1, XCL2*), T cell signaling and maturation (*CD3E, FYB1, THEMIS, TIGIT, IKT, KLRK1*), innate immune response (*GBP5 GBP7, HDC, TLR10*) and other (*CD27, PYHIN1, RNF135*). The mRNA expression of *CD226, CD3D, CD3G, GPB4*, and *UBASH3A* showed no prognostic significance in DFS or OS (Figure 6, A, B, and C). The *CD3D, CD3G*, and *UBASH3A* genes have been shown to contribute to T cell receptor (TCR) synthesis (27-28); *CD226* is involved in lymphocyte adhesion, signaling and cytotoxicity (29), and *GBP4* to be related to interferon-gamma signaling contributing to the innate immunity (30). The *CCRL2* gene mRNA expression emerged as a key feature – it was the only immune gene upregulated in CS-ACC and was purported significant association with poor DFS (HR 1.41, 95% CI 1.02 – 1.94, p=0.036). The *CCRL2* gene codes for the C-C Motif Chemokine Receptor-Like 2, a non-signaling seven-transmembrane domain (7-TMD) receptor related to the atypical chemokine (ACKR) family, however, and its role of this receptor in TME is elusive. ACKRs typically bind chemokines without G protein signaling activation to promote ligand internalization and degradation, however, more importantly, they regulate immune functions by scavenging chemokines from the local environment. Previous studies have demonstrated CCRL2 receptors to act as decoy receptors scavenging chemokines from the TME and their expression to be associated with poor dendritic cell trafficking (31-33). Elevated *CCRL2* expression has been shown in primary neutrophils relative to other immune cell types and further upregulated on neutrophil activation (32-33).

**Figure 6.**
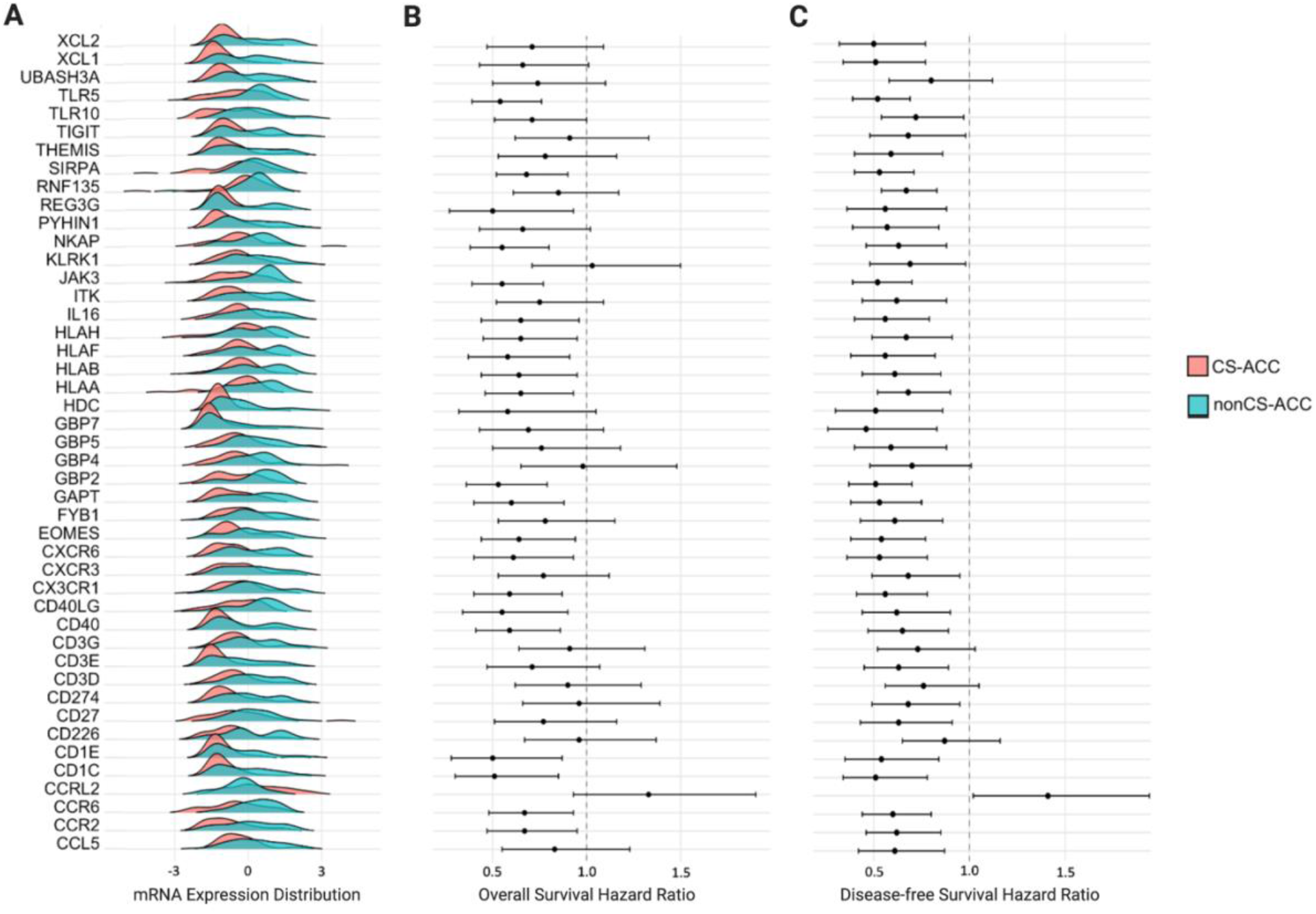
Patient Outcomes Patterns of Immune-related Differentially Expressed Genes (DEGs) in Adrenocortical Carcinoma (ACC). (**A)**. The mRNA expression of 19 immune-related DEGs in CS-ACC and nonCS-ACC tumors. (**B) and (C)**. Significant prognostic indicators for the 19 immune-related DEGs with positive associations for both overall and disease-free survival (OS and DFS) in CS-ACC (HR >1.00, OS and DFS, p<0.05). Univariate Cox regression hazard ratio (HR) and 95% Confidence interval (95% CI). Dot indicates HR and brackets represent 95% CI range.

Genes with mRNA expression found to be significantly associated with OS and DFS (n=19) were compiled to create a composite mRNA expression with immunosuppressive signatures characteristic of CS-ACC tumors. This mRNA expression was made up of *CCR2, CDC1C, CD1E, CD40LG, CD40, CDCR6, EOMES, GAPT, GBP2, HLAA, HLAB, HLAF, HLAH, IL16, JAK3, NKAP, REG3G, SIRPA*, and *TLR5*. The bulk of the genes contributing to the prognostic mRNA signature downregulated in CS-ACC have been shown to code for interactive proteins crucial for T cell-mediated cancer cell killing (34-35). These necessary steps include membrane and intercellular signaling proteins involved in T cell activation (*CDC1C, CD1E*), recruitment (*CCR6, CXCR6, CCR2, IL16*), and tumor recognition (*CDC1C, CD1E, HLAA, HLAB, HLAF, HLAH*) as well as CD8^+^ T cell differentiation (*EOMES*) as illustrated in Figure 7 (35-39). Other gene products, including those of *GBP2, IL16, REG3G*, and *TLR5*, have been shown to contribute to the innate immune response through macrophages activation and enhances phagocytic and oxidative killing (40-46). Signal regulatory protein alpha (*SIRPA*) gene codes for the cell surface receptor for *CD47*. The *SIRPA*-*CD47* has been shown to prevent the maturation of dendritic cells and promote immune tolerance of mature dendritic cells (47-48). Janus kinase (*JAK*) family of tyrosine kinases involved in cytokine receptor-mediated intracellular signal transduction of both the innate and adaptive immune system and mutations of this gene are characteristic of severe combined immunodeficiency (49-50).

**Figure 7.**
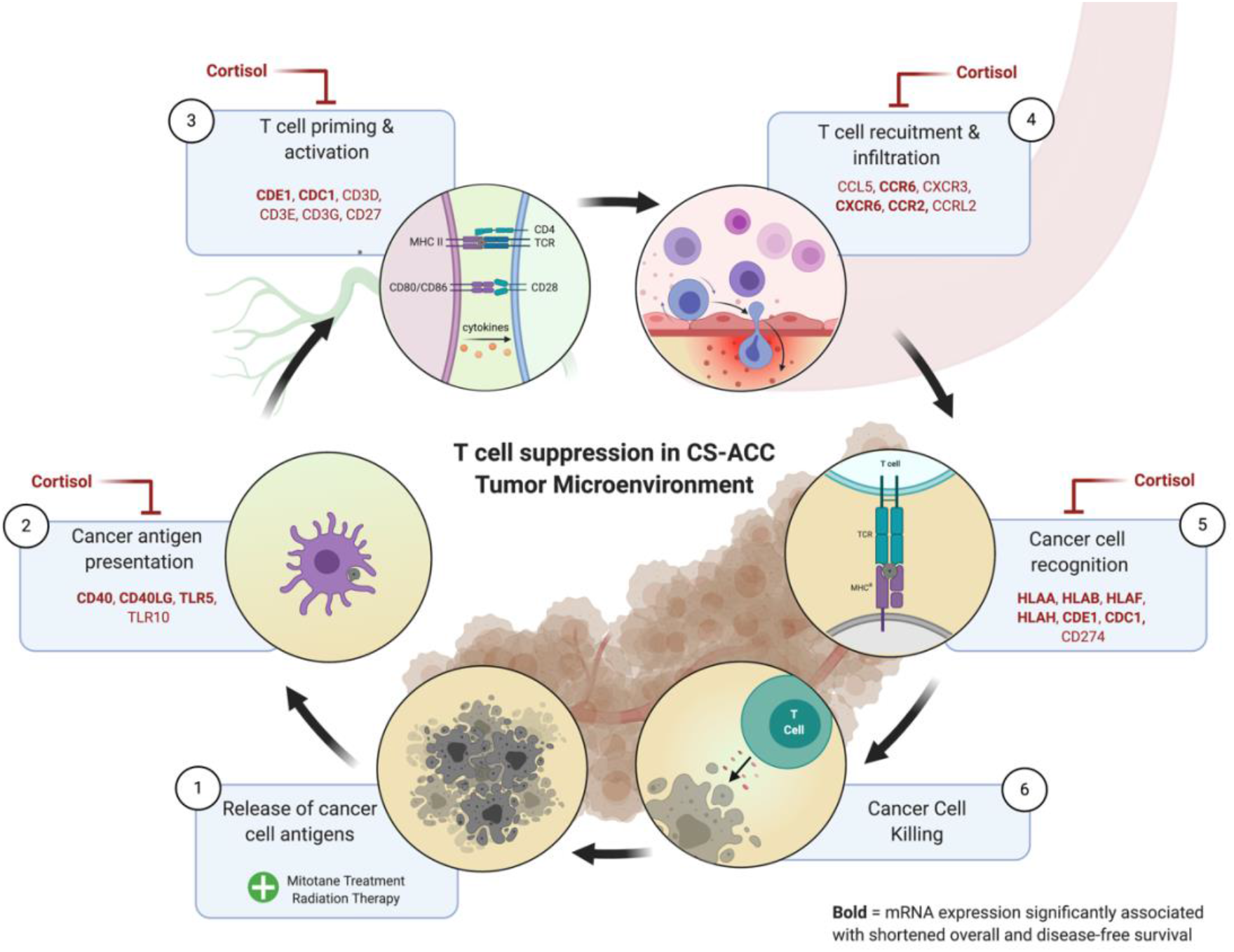
Mechanisms of T Cell Inhibition in Cortisol-Secreting Adrenocortical Carcinoma (CS-ACC). Schematic representation of the steps in cancer cell antigen release (1) and T cell-mediated tumor cell death: (2) tumor cell antigen presentation, (3) T cell priming/activation, (4) T cell recruitment/infiltration, (5) tumor cell recognition (6) T-cell mediated tumor cell death. Genes related to each step in CS-ACC tumors are labeled, with bolded genes signifying their respective prognostic value in overall and disease-free survival (OS and DFS, p<0.05).

The CS-ACC immunogenomic deconvolution revealed an immunosuppressive signature with multiple intercorrelated gene mRNA expression sub-clusters. The HLA subcluster (HLA-A, B, F, H) (r^2^≥0.83) showed strong positive intercorrelations. Furthermore, *CCR2* and *IL16* (r^2^=0.85), as well as *CCR2* and *CXCR6* (r^2^=0.92), showed strong positive correlations (Figure 8A). The strong associations between *CCR2* and *IL16*, which are code for signaling molecules responsible for monocyte recruitment and macrophage activation, respectively, suggest that macrophage tumor infiltration and activation may be highly influential in ACC TME and carry similar gene regulation mechanisms. This would be supported by the TIIC analysis showing decreased monocytes, M1 and M2 macrophages in CS-ACC, and decreased monocyte and M2 macrophage infiltration to associated with shorter OS and DFS, respectively. The strong associations between the HLA gene mRNA expression values would also suggest a decreased MHC class I surface expression, which would result in decreased antigen presentation and T cell activation. Supportively, decreased HLA subcluster mRNA expression was negatively associated with CD8^+^ T cell infiltration.

**Figure 8.**
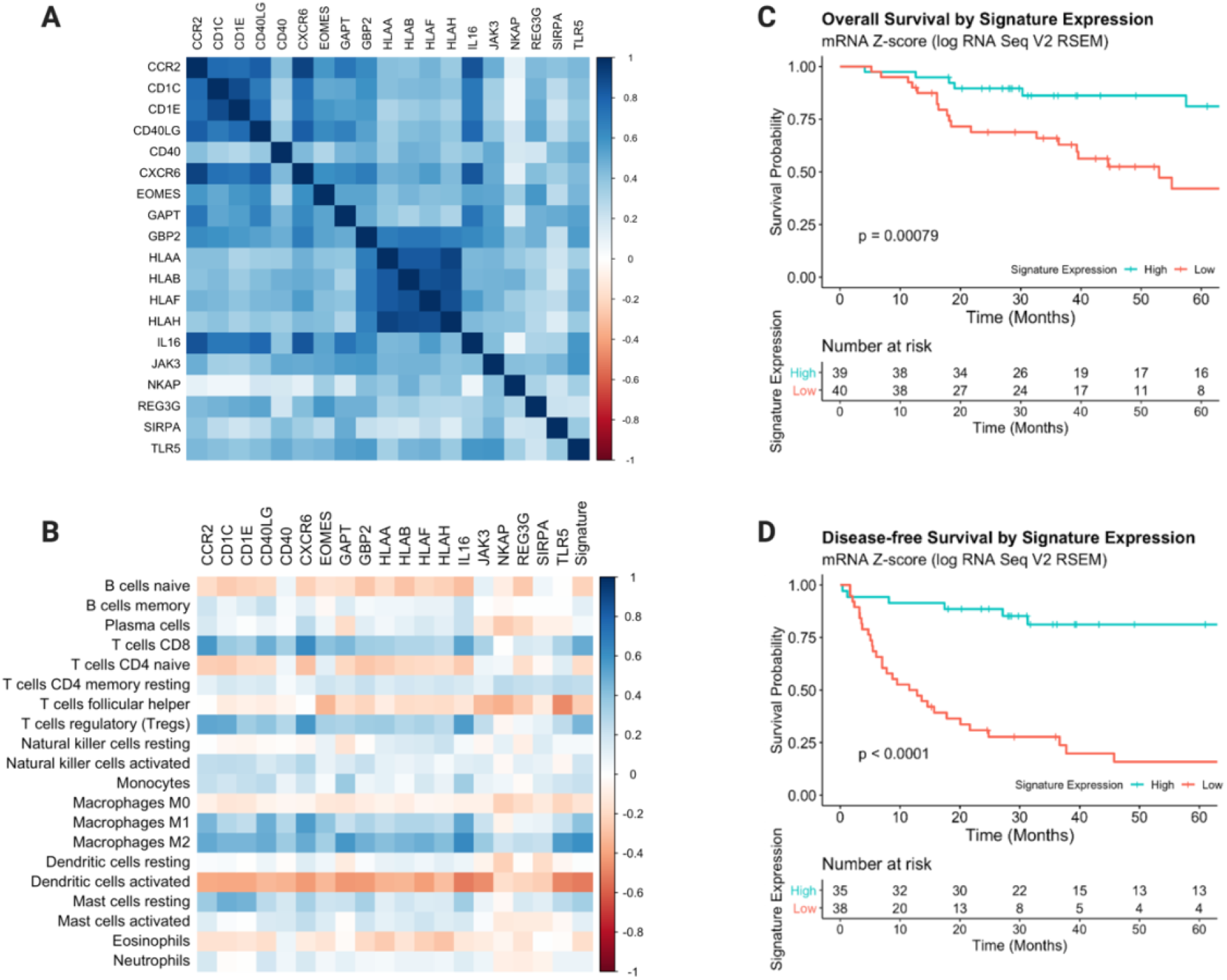
Immunosuppressive Signature of Cortisol-Secreting Adrenocortical Carcinoma (CS-ACC). **(A)**. Heatmap of immunosuppressive signature of CS-ACC with multiple intercorrelated gene mRNA expression sub-clusters. (**B**.) Heatmap of individual gene expression contributing to CS-ACC immunosuppressive signature and significant correlations with tumor-infiltrating immune cell (TIIC) subtypes. (**C)**. Overall survival (OS) comparing low and high expression of the total signature of ACC tumors. (**D)**. Disease-free survival (DFS) comparing low and high expression of the total signature of ACC tumors.

The relationships between the individual gene expression contributing to CS-ACC immunosuppressive signature and the signature as a whole uncovers important correlations with TIICs (Figure 8B). Expression of *CCR2*, a gene primarily expressed in T cells in the recruitment of monocytes, was strongly associated with CD8^+^ T cell (r^2^=0.57), immunosuppressive T regulatory (T_reg_) cells (r^2^=0.52), and M2 macrophage (r^2^=0.51) tumor infiltration. Messenger RNA expression of *CD1C*, a TCR contributory gene, was strongly associated with immunosuppressive T_reg_ infiltration (r^2^=0.50). CD40LG was strongly associated with M2 macrophage infiltration (r^2^=0.50). CXCR6, gene expressed in T cell to mediate further T cell recruitment during inflammation was strongly associated with CD8^+^ T cell (r^2^=0.62), T_reg_ (r^2^=0.57), M1 (r^2^=0.55) and M2 macrophage (r^2^=0.50) infiltration. GAPT expression was associated with increased M2 macrophage infiltration (r^2^=0.57). IL16, a modulator of macrophage and T cell activation, mRNA expression was strongly associated with CD8^+^ T cell (r^2^=0.53), T_reg_ (r^2^=0.56), M1 (r^2^=0.52) and M2 (r^2^=0.64) macrophages and negatively associated with DC_a_ (r^2^=-0.52). TLR5 expression was positively associated with M2 macrophage infiltration (r^2^=-0.49) and T follicular helper cells (r^2^=-0.48). Lastly, the prognostic mRNA expression signature suppressed in CS-ACCs was positively associated with CD8^+^ T cell (r^2^=0.49) and M2 macrophage (r^2^=0.61) infiltration and negatively associated with DC_a_ (r^2^=-0.52) in the ACC TME, suggesting a link between the prognostic gene expression and TIICs.

Comparison analysis of patient outcomes of CS-ACC and nonCS-ACC using univariate Cox regression was further stratified by the expression of the immunosuppressive signature in CS-ACC patients, and revealed that the low expression of the mRNA signature was associated with significantly shorter OS (hazard ratio [HR] 3.98, 95% confidence interval [CI] 1.67 – 9.48, p=0.002) and DFS (HR 7.08; 95% CI 3.07 – 16.33, p<0.001). The 5-year OS for all ACC patients was 81.1% for the high expression group and 42.0% for the low expression group, while the DFS was 81.2% for the high expression group and 15.9% for the low expression group (Figure 8, C and D).

## Discussion

We defined the features of ACC tumor microenvironment and immunosuppressive signatures using multiplatform computational immunogenomic deconvolution methods. Specifically, we noted differences between functional ACC oversecreting cortisol (CS-ACC) and hormonally inactive or non-cortisol-producing hormonally active ACC tumors (nonCS-ACC). Our findings strongly support previous studies where CS-ACC was shown to be associated with poor patient outcomes compared to nonCS-ACC despite similar pathology and stage (13-14). Furthermore, we demonstrated distinct immunogenomic landscape differences between CS-ACC and nonCS-ACC TME while identifying distinct mRNA expression profiles characteristic of CS-ACC TME that were associated with the suppression of immune process genes. The downregulation of many of these genes was associated with poor patient outcomes and differential TIIC profiles. NonCS-ACC tumors demonstrated significantly lower levels of CD8^+^ T cells compared to CS-ACC and the CD8^+^ T cell depletion in the TME was associated with poor prognosis in ACC patients. DC_a_ tumor infiltration was observed to a greater degree in CS-ACC tumors and associated with poor prognosis. Altogether, these findings strongly suggest excess ACC cortisol secretion can exert influence in the TME which may enhance tumor immune escape, potentiate immunotherapeutic failure, and adversely impact patient outcomes.

CS-ACC tumors have long been considered the more aggressive phenotype among ACC tumors. Despite the well-known immunosuppressive role of supra-physiologic glucocorticoid levels, this is the first human study to evaluate the immunogenomic impact of cortisol excess on ACC TME and its clinically relevant impact on patient prognosis. It is well understood that glucocorticoids play a key regulatory role in the cell transcription process and homeostasis. Previous studies have demonstrated major alterations in immune cell genome expression under the treatment of exogenous glucocorticoids (51-52). In our study, we have shown that about 1 in 20 of the genes with mRNA quantification in the TCGA database showed significantly different expression between CS- and nonCS-ACC. Consistent with previous studies, DEGs were primarily related to cellular and metabolic processes and biological regulation, however, a small portion was identified to be directly related to immunological processes. This deductive analysis served as a starting point for our study to further define the potential immunosuppressive role of excess cortisol in the ACC TME.

Cortisol (hydrocortisone) is a corticosteroid with both glucocorticoid and mineralocorticoid activity that is physiologically regulated by the hippocampus-pituitary-adrenal (HPA) axis. CS-ACC tumors escape the HPA negative feedback loop, leading to cortisol concentrations often over 3-fold the upper limit of normal. Recent studies have characterized a variety of mechanisms by which excess cortisol and synthetic cortisol-like therapeutics (prednisone, betamethasone, etc.) impair the immune response and effects of immunotherapy in various cancer types through immune cell deactivation, dampen immune cell recruitment and maturation as well as the induction of apoptosis in lymphocytes (53-54). Supraphysiologic doses of exogenous glucocorticoids are associated with poor immune checkpoint inhibitor (ICI) response, including programmed death receptor-1/ligand-1 monoclonal antibodies and cytotoxic T lymphocyte-associated antigen-4 monoclonal antibodies (53-54). The mechanistic failure of ICIs in the setting of excess glucocorticoids has been mostly attributable to multimodal T cell inhibition and deactivation. Importantly, however, glucocorticoids have been shown to regulate cytokine secretion in T lymphocytes and potentiate the inhibitory capacity of programmed cell death 1 by upregulating its expression on T cells (55-56). In this study, cortisol secretion was associated with a powerful suppression linked to decreased cytotoxic T cell activity in the TME compared to nonCS-ACC. Decreased expression of CD8^+^ T cells promoting chemokines and cell surface proteins, as well as the decreased infiltration of CD8^+^ T cells, were both significant predictors of poor survival in ACC patients. This collage of evidence suggests combating glucocorticoid suppression of CD8^+^ T cell may be a potential avenue for enhancing tumor immunity and immunotherapeutic responses in CS-ACC patients.

Glucocorticoids inhibit macrophage differentiation towards a proinflammatory phenotype by attenuating the induction of pro-inflammatory genes that inhibits their differentiation to an M1 phenotype (57-59). In our study, we observed significant M2 macrophage infiltration to be the predominant macrophage phenotype in all ACC tumors. Significantly fewer activated M1 and M2 macrophages were noted within TME of CS-ACC compared to nonCS-ACC tumors, and the decrease in M2 macrophages was associated with shorter OS and DFS. The decreased M2 macrophage infiltration observed among CS-ACC compared to nonCS-ACC may contribute to the poor prognosis associated with CS-ACC, therefore, increasing macrophage recruitment, maturation, and activation may be an effective future therapeutic endeavor in patients with CS-ACC. Dendritic cells play a key role in immune cell signaling in TME with an emerging role in orchestrating immune activity in the TME. Mature activated dendritic cells are equipped to capture antigens and to produce large numbers of immunogenic MHC-peptide complexes to potentiate T cell immunity. Glucocorticoids have been shown to inhibit the terminal maturation of already differentiated dendritic cells. The overall impact of glucocorticoids on dendritic cells has been summarized as a partial conversion to a monocyte-macrophage phenotype and impaired capacity to reach maturation resulting in decreased T cell stimulation (60-62).

Together, these findings may, in part, explain the elevated levels of DC_a_ cells in CS-ACCs and inefficient antigen presentation. Although our analysis is limited from investigating dendritic cell maturation and efficiency, the accumulation of activated but immature dendritic cells may be a result of cortisol accumulation within the TME due to decreased maturation and peripheral migration. Nonetheless, the strong correlation between DC_a_ in the TME and poor patient prognosis in CS-ACCs certainly justifies further investigation. Stimulation of the glucocorticoid receptor directly associates with NF-κB family proteins to inhibit their transcriptional activity. This results in innate and adaptive immune suppression through the decreased expression of co-stimulatory molecules, cytokines, and chemokines as well as the upregulation of co-inhibitory molecules (63-64). As observed in this study, several downstream NF-κB product genes showed downregulation in CS-ACC (*CCR2, CD40, CD40LG, EOMES, TLR5*).

Our study suggests that the ACC cortisol hypersecretion may exert an influence on TME and can potentially facilitate more aggressive tumor biology and immune resilience. To date, several studies have now highlighted the negative effects of synthetic glucocorticoids on the outcome of immunotherapy. In line with this, patients with CS-ACC were recently reported to experience decreased response to immunotherapy and experience poor patient outcomes compared to nonCS-ACC patients when treated with anti-PD-L1 agent Pembrolizumab and mitotane (23). Studies by Wolkersdörfer et al. suggest the immune escape of ACC may be the consequence of altered Fas/Fas-L system expression and loss of MHC class H and HLA expression in a cell line stimulated to secrete cortisol (65). The decreased expression of HLA-A, B, F, H among CS-ACC and their strong associations with poor patient prognosis observed in this study would further support these findings and may, at least in part, underlie the decreased CD8^+^ T cell response among CS-ACCs.

Future studies will need to clarify the potential mechanisms of immune resistance and immunotherapy failure in CS-ACC. Such insight may empower strategies to reduce the harmful effects of excess cortisol secretion and allow synthetic glucocorticoids to be used to control side effects and symptoms associated with many immunotherapies while preventing them from promoting cancer growth, metastasis, and therapy resistance. In summary, our study characterized a distinct immunogenomic profile associated with CS-ACC compared to nonCS-ACC with a significant prognostic value that may contribute to the poor outcomes associated with CS-ACC, however, in-depth future studies aimed at uncovering the full impact of excess glucocorticoid secretion in TME may provide new immunotherapeutic applications for effective treatment of aggressive and poorly responsive CS-ACC tumors and impact patient survival.

## Methods

### Data Acquisition, Patient Demographics & Tumor Pathology

We utilized the RNA sequencing count table data of Adrenal Cortical Carcinomas (N=92) from The Cancer Genome Atlas (TCGA) Firehose Legacy Cohort (66). Of the 92 patients in the TCGA cohort, 79 (86%) had reported mRNA expression values and were included in the study cohort. The American Joint Commission on Cancer Staging Manual, seventh edition, was used to determine TNM classification. Overall survival (OS) and disease-free survival (DFS) were defined as the time from the date of index operation to the date of death or date of documented disease recurrence or death, respectively. Categorical variables were presented as frequency and percentages and compared using Chi-square or Fisher’s exact test, as appropriate. Continuous variables were reported as median values with interquartile range (IQR) and compared using the Kruskal–Wallis test. OS and DFS were calculated using the Kaplan–Meier method and compared using the log-rank test. Individual and concurrent GRAS components were analyzed using univariant Cox regression methods. Significance was set at a p-value less than 0.05. All statistical analyses were performed using the 1.1.383 R statistics software (R Core Team Vienna, Austria).

### Multiplatform Immunogenomic Deconvolution

*CIBERSORTx* was used to estimate tumor-infiltrating immune subsets (including B cells, CD4^+^T cells, CD8^+^T cells, dendritic cells, macrophages, natural killer cells, neutrophils, etc). *CIBERSORTx* is a computational immunogenomic platform, a publicly available web-based deconvolution program (https://cibersortx.stanford.edu) (67). All significant immune cell and TME DEGs associations were constructed in heatmap format to represent all potential associations. After characterizing the relationships between DEGs and comparing the expression profiles between CS-ACC and nonCS-ACC TMEs. These genes were categorized according to biological function using Panther Gene Classification (68). We then investigated the association of differentially expressed immune process genes with patient outcomes. The Cancer Genome Atlas (TCGA, Firehose Legacy) was accessed through cBioPortal (https://www.cbioportal.org/) (Study ID: 5fe222f5e4b015b63e9d16e8) (69). Gene mRNA expression using z-score relative to all samples (log RNA Seq V2 RSEM) were analyzed in their relation to patient OS and DFS. All survival analyses were performed using the 1.1.383 R statistics software (R Core Team Vienna, Austria).

### Patient Outcomes Analysis

Survival analysis was analyzed by time-to-event Cox regression models controlling for clinical stage and multivariable regression log-rank tests were used to assess overall survival (OS) and disease-free survival (DFS) hazard ratio (HR) statistical significance. Kaplan–Meier method was used to compare the OS of ACC patients according to mRNA expression signature profiles. Gene expression signatures were compiles by normalizing the sum of the gene mRNA Z-scores (log RNA Seq V2 RSEM) relative to the median on a scale of −5 to 5. Patients with positive cumulative normalized expression levels (≥0.00) were assigned to the high signature expression group and those with negative cumulative normalized expression levels (<0.00) were assigned to the low signature expression group. Kaplan–Meier method OS analysis was compared using the univariate log-rank test. Significance was set at a p*-*value lower than 0.05 for all analysis types. All statistical comparisons were performed using the 1.1.383 R statistics software (R Core Team Vienna, Austria).

## Data Availability

The Cancer Genome Atlas (TCGA, Firehose Legacy) was accessed through cBioPortal (https://www.cbioportal.org/) (Study ID: 5fe222f5e4b015b63e9d16e8) (69).

## Statistics

All survival analyses and statistical comparisons were performed using the 1.1.383 R statistics software (R Core Team Vienna, Austria).

## Author contributions

NB, JJB, JCR and WKR designed the study. NB, JJB, DNH, SRK performed genomic and correlative clinical outcomes analyses. NB and JJB generated figures. NB and JJB wrote the manuscript with input from all authors.

## Acknowledgments

None

