## Supplemental Table 1 for "Multiplatform Integrative Analyses of Immunosuppressive Signatures in Cortisol-secreting Adrenocortical Carcinoma"

**Supplemental material**

**Supplemental Table 1. Patient Outcomes by Differential mRNA Expression of Immune-related Genes in Adrenocortical Carcinoma.** The comparative distribution of DEGs by cortisol secretion and the impact of DEGs on OS and DFS.

| | Overall Survival | | Disease-Free Survival | |
|---|---|---|---|---|
| | Hazard Ratio [95% Confidence Interval] | p-value | Hazard Ratio [95% Confidence Interval] | p-value |
| *CCR2* | 0.67 [0.47 – 0.95] | 0.026 | 0.62 [0.46 – 0.85] | 0.003 |
| *CCL6* | 0.67 [0.48 – 0.93] | 0.018 | 0.60 [0.44 – 0.80] | 0.001 |
| *CD1C* | 0.51 [0.30 – 0.85] | 0.010 | 0.51 [0.34 – 0.78] | 0.002 |
| *CD1E* | 0.50 [0.28 – 0.87] | 0.014 | 0.54 [0.35 – 0.84] | 0.006 |
| *CD40LG* | 0.55 [0.34 – 0.90] | 0.016 | 0.62 [0.44 – 0.90] | 0.011 |
| *CD40* | 0.59 [0.41 – 0.86] | 0.006 | 0.65 [0.47 – 0.89] | 0.007 |
| *CXCR6* | 0.61 [0.40 – 0.93] | 0.023 | 0.61 [0.42 – 0.87] | 0.007 |
| *EOMES* | 0.64 [0.44 – 0.94] | 0.022 | 0.54 [0.38 – 0.77] | 0.001 |
| *GAPT* | 0.60 [0.40 – 0.88] | 0.009 | 0.53 [0.38 – 0.75] | <0.001 |
| *GBP2* | 0.53 [0.36 – 0.79] | 0.007 | 0.51 [0.37 – 0.70] | <0.001 |
| *HLAA* | 0.65 [0.46 – 0.93] | 0.017 | 0.68 [0.52 – 0.90] | 0.006 |
| *HLAB* | 0.64 [0.44 – 0.95] | 0.025 | 0.61 [0.44 – 0.85] | 0.003 |
| *HLAF* | 0.58 [0.37 - 0.91] | 0.017 | 0.56 [0.38 – 0.82] | 0.003 |
| *HLAH* | 0.65 [0.45 – 0.95] | 0.026 | 0.67 [0.49 – 0.91] | 0.010 |
| *IL16* | 0.65 [0.44 – 0.96] | 0.031 | 0.56 [0.40 – 0.88] | 0.001 |
| *JAK3* | 0.55 [0.39 – 0.77] | 0.001 | 0.52 [0.39 – 0.70] | <0.001 |
| *NKAP* | 0.55 [0.38 – 0.80] | 0.002 | 0.63 [0.46 – 0.88] | 0.006 |
| *REG3G* | 0.50 [0.27 – 0.93] | 0.028 | 0.56 [0.36 – 0.88] | 0.011 |
| *SIRPA* | 0.68 [0.52 – 0.90] | 0.007 | 0.53 [0.40 – 0.71] | <0.001 |
